# Hypoxic Modulation of Dental Pulp Stem Cells Viability: an *In Vitro* Study

**DOI:** 10.64898/2026.06.09.731107

**Authors:** Francesco Torelli

## Abstract

**Introduction:** To investigate the effects of chronic hypoxic exposure simulating heavy-industrial environments on the viability, metabolic activity, stemness preservation, and osteogenic differentiation potential of human dental pulp stem cells (hDPSCs).

**Methods:** Commercially available hDPSCs were cultured under controlled oxygen tensions representing surface atmospheric conditions (21% O_2_), moderate hypoxia (10% O_2_), deep hypoxia (5% O_2_), and severe hypoxia (1% O_2_). Cells were maintained for 1, 3, and 7 days. Cell viability was evaluated using MTT and Live/Dead assays. Reactive oxygen species (ROS) accumulation, mitochondrial membrane potential, and apoptosis were assessed using fluorescent probes and Annexin V/PI staining. Stemness marker expression (SOX2, OCT4, NANOG) and osteogenic markers (RUNX2, ALP, OCN) were analyzed via RT-qPCR.

**Results:** Moderate hypoxia (10% O_2_) promoted transient increases in stemness marker expression and preserved metabolic activity. Severe hypoxia (1% O_2_) significantly reduced cell viability, increased ROS accumulation, disrupted mitochondrial integrity, and elevated apoptotic cell populations after prolonged exposure (p < 0.05). Osteogenic differentiation markers were significantly downregulated under severe hypoxic conditions.

**Conclusions:** Industrial hypoxic environments critically influence pulpal stem cell physiology and regenerative potential. While moderate oxygen reduction may transiently preserve stemness characteristics, chronic severe hypoxia impairs viability and osteogenic functionality. Chronic low-oxygen occupational environments may alter endogenous dental regenerative mechanisms and influence oral tissue healing responses.

## Introduction

Human dental pulp stem cells (hDPSCs) are a mesenchymal stem cell population with significant potential for regenerative dentistry and craniofacial tissue engineering due to their proliferative capacity, multilineage differentiation ability, and accessibility from dental tissues [1–3]. Their application has been extensively investigated in dentin–pulp regeneration, bone tissue engineering, and regenerative oral medicine [2–6].

Stem cell behavior is strongly influenced by the physicochemical characteristics of the surrounding microenvironment, among which oxygen tension plays a critical regulatory role [7,8]. Oxygen availability modulates cellular metabolism, proliferation, differentiation, survival, and self-renewal through pathways primarily mediated by hypoxia-inducible factors (HIFs) [8-10]. Importantly, physiological stem cell niches typically exist under oxygen concentrations substantially lower than atmospheric conditions, commonly ranging between 1% and 7% O_2_ [7].

Conventional *in vitro* culture is generally performed under atmospheric oxygen concentrations approximating 21% O_2_, despite evidence that these levels exceed those physiologically encountered within many mesenchymal tissues, including dental pulp [7,8,11]. Due to its confined vascular architecture and metabolic activity, the dental pulp is particularly sensitive to fluctuations in oxygen availability [8,12]. Consequently, standard atmospheric incubation may inadequately reproduce the native biological environment of hDPSCs [7,12].

Previous studies suggest that moderate hypoxia may preserve stemness characteristics, promote glycolytic metabolism, and reduce oxidative stress in mesenchymal stem cells [7,8,10]. In dental-derived stem cells specifically, controlled hypoxic exposure has been associated with maintenance of undifferentiated phenotypes and clonogenic potential [7]. Conversely, prolonged or severe hypoxia may induce mitochondrial dysfunction, reactive oxygen species (ROS) accumulation, apoptosis, and suppression of osteogenic differentiation pathways, thereby compromising regenerative capacity [13–15].

Although hypoxia-mediated modulation of mesenchymal stem cells has been increasingly investigated, the transition between adaptive physiological hypoxia and detrimental oxygen deprivation remains insufficiently characterized in hDPSCs [14,15]. This question is relevant not only for optimizing stem-cell-based regenerative strategies and *in vitro* culture methodologies, but also for understanding how reduced oxygen availability may influence endogenous pulpal healing responses [7,12,16].

Therefore, the aim of the present study was to evaluate the effects of simulated hypoxic environmental conditions on the viability, oxidative stress response, apoptosis, stemness maintenance, and osteogenic differentiation potential of commercially available human dental pulp stem cells *in vitro*. We hypothesized that moderate hypoxia would transiently preserve stemness-related characteristics and metabolic activity, whereas severe chronic hypoxia would induce oxidative stress, mitochondrial dysfunction, apoptosis, and impaired osteogenic differentiation.

## Materials & Methods

### Study Design

An *in vitro* experimental study was designed to evaluate the effects of controlled hypoxic conditions on human dental pulp stem cell (hDPSC) viability, oxidative stress response, apoptosis, stemness maintenance, and osteogenic differentiation. Oxygen tensions were selected to reproduce progressively hypoxic environments ranging from conventional atmospheric culture conditions to severe hypoxia [17–19]. All experiments were independently repeated three times.

### Cell Culture

Commercially available human dental pulp stem cells (hDPSCs; ATCC-certified) were acquired from Lonza. Cells at passage 3 or 4 were selected to minimize phenotypic drift [20–22]. Cells were cultured in α-minimum essential medium supplemented with 10% fetal bovine serum and 1% penicillin/streptomycin under standard humidified conditions (37°C, 5% CO_2_). Culture medium was replaced every 48 h.

### Experimental Groups and Hypoxic Exposure

Cells were divided into four experimental groups according to oxygen concentration (**Table 1**).

**Table 1:** Experimental groups design.

| Group | Oxygen Level [21,23] |
| --- | --- |
| CTRL | 21% O<sub>2</sub> |
| MOD | 10% O<sub>2</sub> |
| DEEP | 5 % O<sub>2</sub> |
| CRIT | 1% O<sub>2</sub> |

Hypoxic exposure was performed using oxygen-regulated incubators under controlled temperature, humidity, and CO_2_ conditions [18,21,23]. Oxygen concentrations were continuously stabilized throughout the experimental period. Exposure durations were established as Day 1, Day 3, and Day 7. The 21% O_2_ group served as conventional atmospheric control conditions [24].

### Cell Viability Assessment

Cell metabolic activity was evaluated using MTT assay [17,18]. Cells were seeded into 96-well plates (1 × 10^4^ cells/well) and exposed to experimental oxygen conditions for the designated timepoints. Absorbance was measured at 570 nm, and results were normalized relative to the CTRL group. Cell viability was further assessed using Live/Dead fluorescence staining with calcein-AM and ethidium homodimer-1 [23]. Fluorescence images were acquired using inverted fluorescence microscopy, and quantitative analysis was performed using ImageJ software.

### Oxidative Stress and Mitochondrial Function

Intracellular reactive oxygen species (ROS) accumulation was evaluated using DCFH-DA fluorescent staining [23,25]. Fluorescence intensity was quantified using ImageJ and normalized to cell number. Mitochondrial membrane potential was analyzed using JC-1 staining [26]. Healthy mitochondria were identified by red fluorescence emission, whereas depolarized mitochondria exhibited green fluorescence. Mitochondrial alterations were quantified using the red/green fluorescence ratio.

### Apoptosis Analysis

Apoptotic cell populations were evaluated using Annexin V-FITC/propidium iodide flow cytometry [25,26]. Following hypoxic exposure, cells were collected, stained according to manufacturer instructions, and analyzed using flow cytometry. Cell populations were categorized as viable, early apoptotic, late apoptotic, or necrotic. Data were analyzed using FlowJo software.

### Stemness Gene Expression

Total RNA extraction, cDNA synthesis, and RT-qPCR were performed according to standard protocols. Stemness-related genes including SOX2, OCT4, and NANOG were analyzed [18]. Gene expression levels were normalized to GAPDH using the 2^-ΔΔCt method [27].

### Osteogenic Differentiation

Following hypoxic exposure, cells were cultured in osteogenic differentiation medium for 14 days [24]. Osteogenic differentiation was evaluated through RT-qPCR analysis of alkaline phosphatase (ALP), runt-related transcription factor 2 (RUNX2), and osteocalcin (OCN) [23]. Gene expression levels were normalized to GAPDH. Extracellular mineralization was assessed using Alizarin Red S staining. Quantitative mineralization analysis was performed spectrophotometrically following dye extraction.

### Statistical Analysis

Quantitative data were expressed as mean ± standard deviation (SD). Statistical analysis was performed using RStudio. Comparisons among groups were conducted using one-way or two-way ANOVA followed by Tukey’s post hoc test [28]. Statistical significance was established at p < 0.05.

## Results

### Cell Viability and Metabolic Activity

Hypoxic exposure significantly influenced hDPSC metabolic activity in an oxygen-dependent manner (**Figure 1**). At Day 1, cells cultured under moderate hypoxia (MOD; 10% O_2_) showed metabolic activity comparable to CTRL conditions, whereas DEEP (5% O_2_) and CRIT (1% O_2_) groups demonstrated mild reductions. By Day 3, MOD cultures maintained significantly higher metabolic activity compared with CRIT (p < 0.05), while DEEP showed intermediate values. At Day 7, severe hypoxia resulted in a marked reduction in metabolic activity. The CRIT group demonstrated significantly lower MTT values compared with all other groups (p < 0.001), whereas MOD maintained viability levels comparable to CTRL conditions. DEEP cultures exhibited moderate but significant reductions relative to CTRL (p < 0.05).

**Figure 1.**
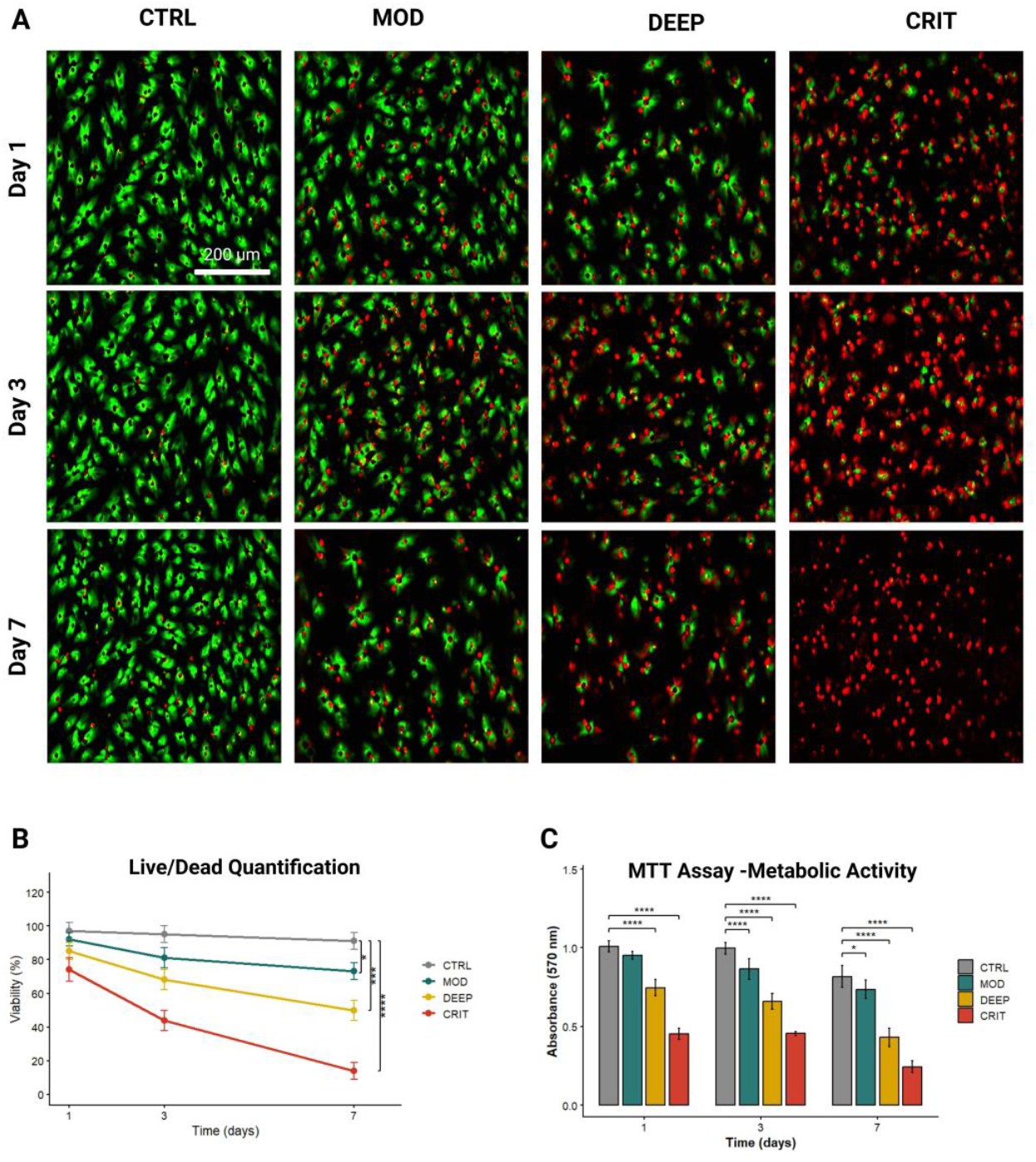
Cell viability and metabolic activity of human dental pulp stem cells (hDPSCs) cultured under simulated hypoxic conditions. A) Representative Live/Dead fluorescence microscopy images obtained at Day 1, Day 3 and Day 7. Viable cells are shown in green (calcein-AM) and dead cells in red (ethidium homodimer). Increased cell death and reduced cellular density were observed under severe hypoxic exposure. Scale bar = 200 μm. B) Quantitative analysis of cell viability percentages over time derived from Live/Dead assays. Chronic severe hypoxia resulted in progressive viability decline compared with normoxic controls. C) MTT assay quantification of metabolic activity following exposure to normoxic (21% O_2_) and hypoxic conditions (10%, 5%, and 1% O_2_) for 1, 3, and 7 days. Moderate hypoxia (10% O_2_) maintained metabolic activity close to control levels, whereas severe hypoxia (1% O_2_) induced a significant reduction over time. Data are presented as mean ± SD (n = 6). Statistical significance: *p < 0.05, **p < 0.01, ***p < 0.001, **** p < 0.0001 versus CTRL.

Live/Dead staining confirmed these findings. CTRL and MOD groups maintained high viable cell density throughout the experimental period, whereas DEEP and particularly CRIT cultures showed progressive reductions in cell density and increased non-viable cell staining after prolonged exposure. Quantitative analysis demonstrated significantly lower viability percentages in CRIT compared with CTRL and MOD groups at Day 3 and Day 7 (p < 0.001).

### Oxidative Stress and Mitochondrial Dysfunction

Intracellular ROS accumulation progressively increased with decreasing oxygen concentration (**Figure 2A,C**). ROS levels remained relatively stable in CTRL and MOD groups, whereas CRIT cultures demonstrated significantly increased fluorescence intensity beginning at Day 1 (p < 0.05) (**Table 2**). By Day 7, CRIT cells exhibited the highest ROS production among all groups (p < 0.001).

**Table 2:** Significance of ROS oxidative stress quantification comparison, as in graph in Figure 2C. * p < 0,05; ** p < 0,01; *** p < 0,001; **** p < 0,0001

| Oxidative Stress Group Comparisons |  |
| --- | --- |
| Group Comparison | Significance |
| CTRL Day 1 vs MOD Day 1 | ** |
| CTRL Day 1 vs DEEP Day 1 | **** |
| CTRL Day 1 vs CRIT Day 1 | **** |
| MOD Day 1 vs DEEP Day 1 | **** |
| MOD Day 1 vs CRIT Day 1 | **** |
| DEEP Day 1 vs CRIT Day 1 | **** |
| CTRL Day 3 vs MOD Day 3 | ** |
| CTRL Day 3 vs DEEP Day 3 | **** |
| CTRL Day 3 vs CRIT Day 3 | **** |
| MOD Day 3 vs DEEP Day 3 | **** |
| MOD Day 3 vs CRIT Day 3 | **** |
| DEEP Day 3 vs CRIT Day 3 | **** |
| CTRL Day 7 vs MOD Day 7 | **** |
| CTRL Day 7 vs DEEP Day 7 | **** |
| CTRL Day 7 vs CRIT Day 7 | **** |
| MOD Day 7 vs DEEP Day 7 | **** |
| MOD Day 7 vs CRIT Day 7 | **** |
| DEEP Day 7 vs CRIT Day 7 | **** |
| CTRL Day 1 vs CTRL Day 3 | **** |
| CTRL Day 1 vs CTRL Day 7 | **** |
| CTRL Day 3 vs CTRL Day 7 | **** |
| MOD Day 1 vs MOD Day 3 | **** |
| MOD Day 1 vs MOD Day 7 | **** |
| MOD Day 3 vs MOD Day 7 | **** |
| DEEP Day 1 vs DEEP Day 3 | **** |
| DEEP Day 1 vs DEEP Day 7 | **** |
| DEEP Day 3 vs DEEP Day 7 | **** |
| CRIT Day 1 vs CRIT Day 3 | **** |
| CRIT Day 1 vs CRIT Day 7 | **** |
| CRIT Day 3 vs CRIT Day 7 | **** |

**Figure 2.**
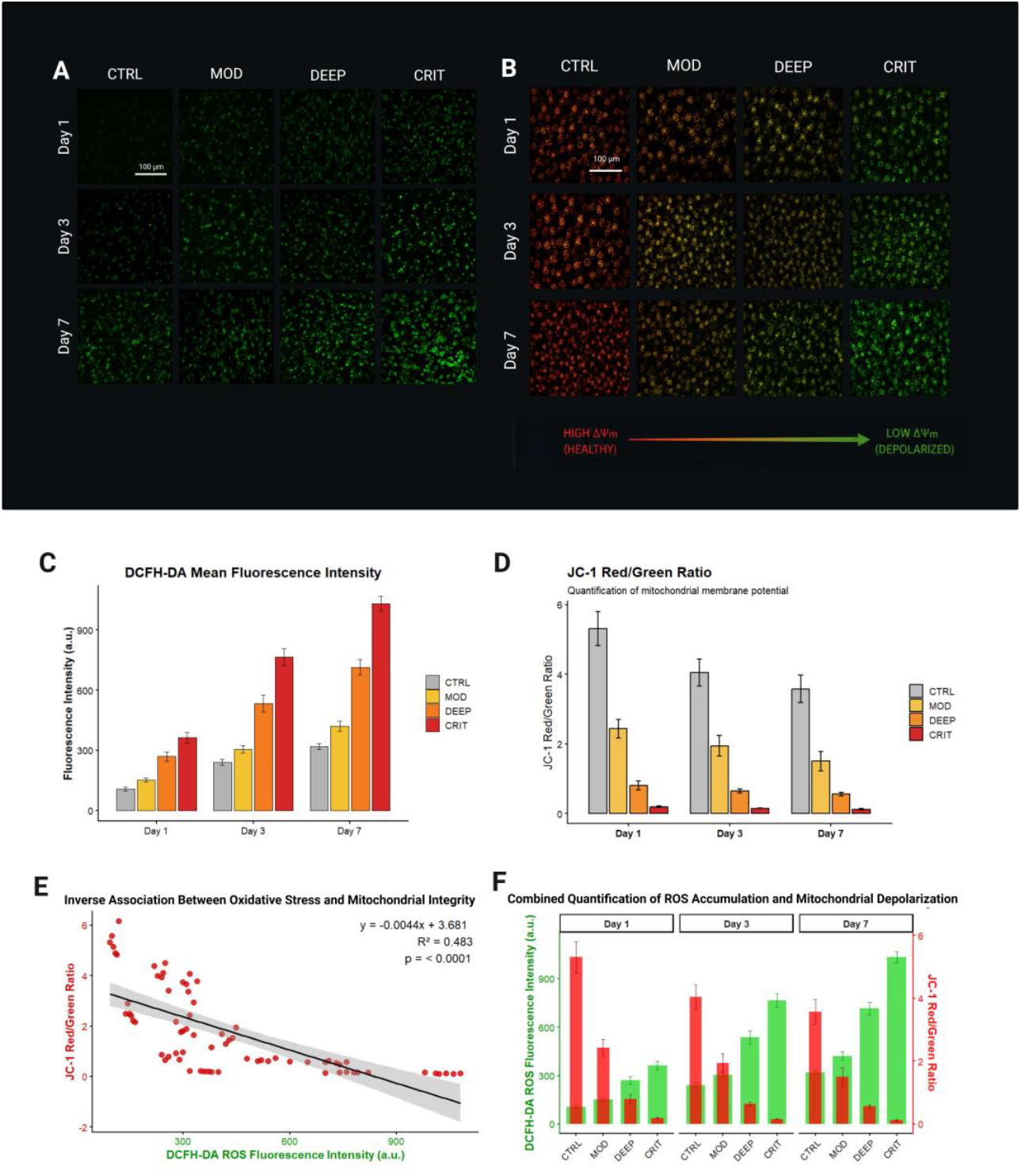
Oxidative stress accumulation and mitochondrial dysfunction in human dental pulp stem cells (hDPSCs) exposed to hypoxic environments. A) Representative DCFH-DA fluorescence micrographs of intracellular reactive oxygen species (ROS) production in hDPSCs cultured under normoxic control conditions (21% O_2_) and hypoxia (10%, 5%, and 1% O_2_) for 1, 3, and 7 days. Progressive ROS accumulation was observed with decreasing oxygen tension and prolonged exposure duration, with the highest fluorescence intensity detected in the SUB-CRIT (1% O_2_) group at Day 7. B) JC-1 staining analysis of mitochondrial membrane potential (ΔΨm). Healthy polarized mitochondria exhibited predominant red fluorescence (JC-1 aggregates), whereas depolarized mitochondria displayed increased green fluorescence (JC-1 monomers). Severe hypoxia induced progressive mitochondrial depolarization, particularly in the SUB-CRIT group. C) Quantitative analysis of DCFH-DA. D) Quantification of JC-1. E) Inverse correlation analysis between intracellular ROS accumulation (DCFH-DA fluorescence intensity) and mitochondrial membrane potential (JC-1 red/green ratio) across all experimental groups and timepoints. Increasing ROS levels were associated with progressive mitochondrial depolarization under severe hypoxic exposure. Pearson correlation analysis was performed to assess the association between ROS accumulation and mitochondrial membrane potential. F) Quantification of oxidative stress and mitochondrial dysfunction across experimental groups, demonstrating increased ROS production and reduced JC-1 red/green fluorescence ratios under severe hypoxic conditions. Data are presented as mean ± SD (n = 6). Statistical significance: *p < 0.05, **p < 0.01, ***p < 0.001. Scale bars = 100 μm.

JC-1 staining demonstrated progressive mitochondrial dysfunction under severe hypoxia (**Figures 2B,D**). CTRL and MOD cultures maintained preserved mitochondrial membrane polarization throughout the study period. In contrast, DEEP cultures showed gradual reductions in mitochondrial membrane potential beginning at Day 3, while CRIT cells exhibited pronounced mitochondrial depolarization at Day 7, with significantly lower red/green fluorescence ratios compared with all other groups (p < 0.001) (**Table 3**).

**Table 3:** Significance of JC-1 mitochondrial membrane depolarization quantification comparison, as in graph in Figure 2D. * p < 0,05; ** p < 0,01; *** p < 0,001; **** p < 0,0001. Only significant comparisons are shown.

| Mitochondrial Membrane Depolarization (JC-1) Group Comparison<sup>a</sup> and Significance |  |
| --- | --- |
| Group Comparison | Significance |
| CTRL Day 1 vs MOD Day 1 | **** |
| CTRL Day 1 vs DEEP Day 1 | **** |
| CTRL Day 1 vs CRIT Day 1 | **** |
| MOD Day 1 vs DEEP Day 1 | **** |
| MOD Day 1 vs CRIT Day 1 | **** |
| DEEP Day 1 vs CRIT Day 1 | *** |
| CTRL Day 3 vs MOD Day 3 | **** |
| CTRL Day 3 vs DEEP Day 3 | **** |
| CTRL Day 3 vs CRIT Day 3 | **** |
| MOD Day 3 vs DEEP Day 3 | **** |
| MOD Day 3 vs CRIT Day 3 | **** |
| DEEP Day 3 vs CRIT Day 3 | ** |
| CTRL Day 7 vs MOD Day 7 | **** |
| CTRL Day 7 vs DEEP Day 7 | **** |
| CTRL Day 7 vs CRIT Day 7 | **** |
| MOD Day 7 vs DEEP Day 7 | **** |
| MOD Day 7 vs CRIT Day 7 | **** |
| DEEP Day 7 vs CRIT Day 7 | * |
| CTRL Day 1 vs CTRL Day 3 | **** |
| CTRL Day 1 vs CTRL Day 7 | **** |
| CTRL Day 3 vs CTRL Day 7 | ** |
| MOD Day 1 vs MOD Day 3 | ** |
| MOD Day 1 vs MOD Day 7 | **** |
| MOD Day 3 vs MOD Day 7 | * |

### Apoptosis Analysis

Flow cytometric analysis revealed oxygen-dependent modulation of apoptotic cell populations (**Figure 3**). Only minor differences were observed at Day 1; however, prolonged hypoxic exposure significantly altered cellular survival profiles. At Day 7, the CRIT group demonstrated marked increases in both early and late apoptotic populations compared with CTRL conditions (p < 0.001) (**Table 4**). Necrotic cell percentages were also significantly elevated under severe hypoxia. Conversely, MOD cultures maintained apoptosis levels comparable to atmospheric conditions, while DEEP cultures demonstrated intermediate responses.

**Table 4:** Significance of early and late apoptosis quantification comparison, as in graph in Figure 3B-C. * p < 0,05; ** p < 0,01; *** p < 0,001; **** p < 0,0001. Only significant comparisons are shown.

| Early Apoptosis Group Comparisons and Significance |  |
| --- | --- |
| Group Comparison | Significance |
| CTRL Day 1 vs MOD Day 1 | ** |
| CTRL Day 1 vs DEEP Day 1 | *** |
| CTRL Day 1 vs CRIT Day 1 | *** |
| MOD Day 1 vs DEEP Day 1 | *** |
| MOD Day 1 vs CRIT Day 1 | *** |
| DEEP Day 1 vs CRIT Day 1 | *** |
| CTRL Day 3 vs MOD Day 3 | *** |
| CTRL Day 3 vs DEEP Day 3 | *** |
| CTRL Day 3 vs CRIT Day 3 | *** |
| MOD Day 3 vs DEEP Day 3 | *** |
| MOD Day 3 vs CRIT Day 3 | *** |
| DEEP Day 3 vs CRIT Day 3 | *** |
| CTRL Day 7 vs MOD Day 7 | *** |
| CTRL Day 7 vs DEEP Day 7 | *** |
| CTRL Day 7 vs CRIT Day 7 | *** |
| MOD Day 7 vs DEEP Day 7 | *** |
| MOD Day 7 vs CRIT Day 7 | *** |
| DEEP Day 7 vs CRIT Day 7 | *** |
| CTRL Day 1 vs CTRL Day 7 | *** |
| CTRL Day 3 vs CTRL Day 7 | * |
| MOD Day 1 vs MOD Day 3 | *** |
| MOD Day 1 vs MOD Day 7 | *** |
| MOD Day 3 vs MOD Day 7 | *** |
| DEEP Day 1 vs DEEP Day 3 | *** |
| DEEP Day 1 vs DEEP Day 7 | *** |
| DEEP Day 3 vs DEEP Day 7 | * |
| CRIT Day 1 vs CRIT Day 3 | *** |
| CRIT Day 1 vs CRIT Day 7 | *** |
| CRIT Day 3 vs CRIT Day 7 | *** |
| Early Apoptosis Group Comparisons and Significance |  |
| CTRL Day 1 vs MOD Day 1 | ** |
| CTRL Day 1 vs DEEP Day 1 | *** |
| CTRL Day 1 vs CRIT Day 1 | *** |
| MOD Day 1 vs DEEP Day 1 | *** |
| MOD Day 1 vs CRIT Day 1 | *** |
| DEEP Day 1 vs CRIT Day 1 | *** |
| CTRL Day 3 vs MOD Day 3 | *** |
| CTRL Day 3 vs DEEP Day 3 | *** |
| CTRL Day 3 vs CRIT Day 3 | *** |
| MOD Day 3 vs DEEP Day 3 | *** |
| MOD Day 3 vs CRIT Day 3 | *** |
| DEEP Day 3 vs CRIT Day 3 | *** |
| CTRL Day 7 vs MOD Day 7 | *** |
| CTRL Day 7 vs DEEP Day 7 | *** |
| CTRL Day 7 vs CRIT Day 7 | *** |
| MOD Day 7 vs DEEP Day 7 | *** |
| MOD Day 7 vs CRIT Day 7 | *** |
| DEEP Day 7 vs CRIT Day 7 | *** |
| CTRL Day 1 vs CTRL Day 7 | *** |
| CTRL Day 3 vs CTRL Day 7 | *** |
| CTRL Day 1 vs CTRL Day 3 | * |
| CTRL Day 1 vs CTRL Day 7 | *** |
| CTRL Day 3 vs CTRL Day 7 | *** |
| MOD Day 1 vs MOD Day 3 | *** |
| MOD Day 1 vs MOD Day 7 | *** |
| MOD Day 3 vs MOD Day 7 | *** |
| DEEP Day 1 vs DEEP Day 3 | *** |
| DEEP Day 1 vs DEEP Day 7 | *** |
| DEEP Day 3 vs DEEP Day 7 | *** |
| CRIT Day 1 vs CRIT Day 3 | *** |
| CRIT Day 1 vs CRIT Day 7 | *** |
| CRIT Day 3 vs CRIT Day 7 | *** |

**Figure 3.**
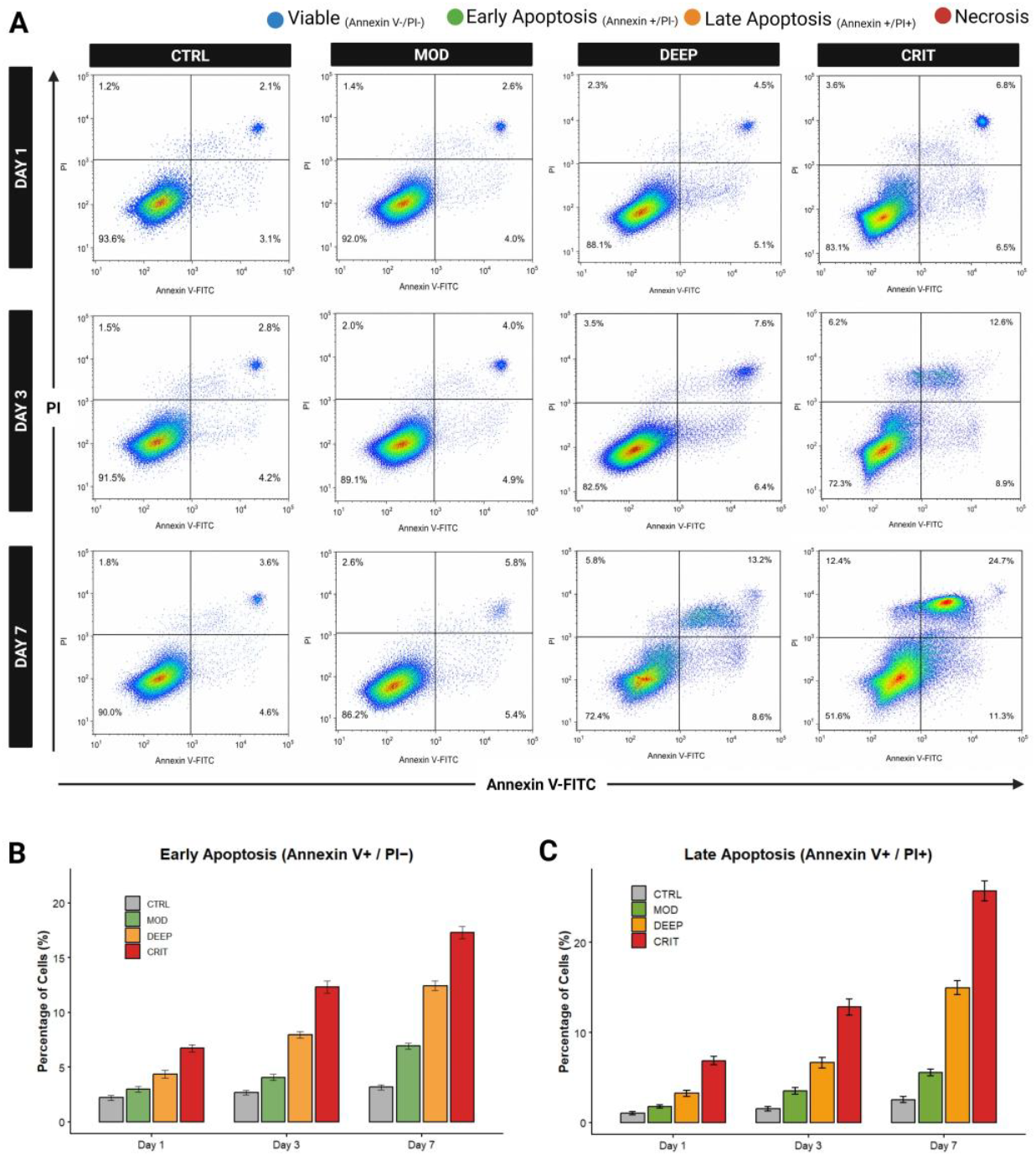
Apoptosis analysis of human dental pulp stem cells (hDPSCs) under hypoxic conditions. A) Representative Annexin V-FITC/PI flow cytometry dot plots of hDPSCs cultured under normoxic control conditions (21% O_2_), moderate hypoxia (10% O_2_), deep hypoxia (5% O_2_), and severe hypoxia (1% O_2_) for 1, 3, and 7 days. Cell populations were classified as viable (Annexin V−/PI−), early apoptotic (Annexin V+/PI−), late apoptotic (Annexin V+/PI+), or necrotic (Annexin V−/PI+). Progressive increases in apoptotic cell populations were observed with decreasing oxygen tension and prolonged exposure duration. B) Quantification of early apoptotic cells (Annexin V+/PI−) across experimental groups and timepoints. Moderate increases were detected under 5% O_2_ conditions, while severe hypoxia (1% O_2_) induced a marked elevation in early apoptosis, particularly at Day 7. C) Quantification of late apoptotic cells (Annexin V+/PI+) demonstrating significant accumulation of late-stage apoptotic populations under severe hypoxic exposure. Data are presented as mean ± standard deviation (SD) (n = 6). Statistical significance was determined using one-way ANOVA followed by Tukey’s post hoc test (*p < 0.05, **p < 0.01, ***p < 0.001 versus CTRL 21% O_2_).

### Stemness Marker Expression

RT-qPCR analysis demonstrated differential modulation of stemness-related genes according to oxygen tension (**Figure 4A**). At Day 3, MOD cultures showed significant upregulation of SOX2, OCT4, and NANOG expression compared with CTRL conditions (p < 0.05), with the strongest effect observed for SOX2. DEEP cultures exhibited modest transient increases at earlier timepoints, followed by progressive decline. In contrast, prolonged severe hypoxia resulted in significant downregulation of all analyzed stemness markers at Day 7 in the CRIT group (p < 0.001).

**Figure 4.**
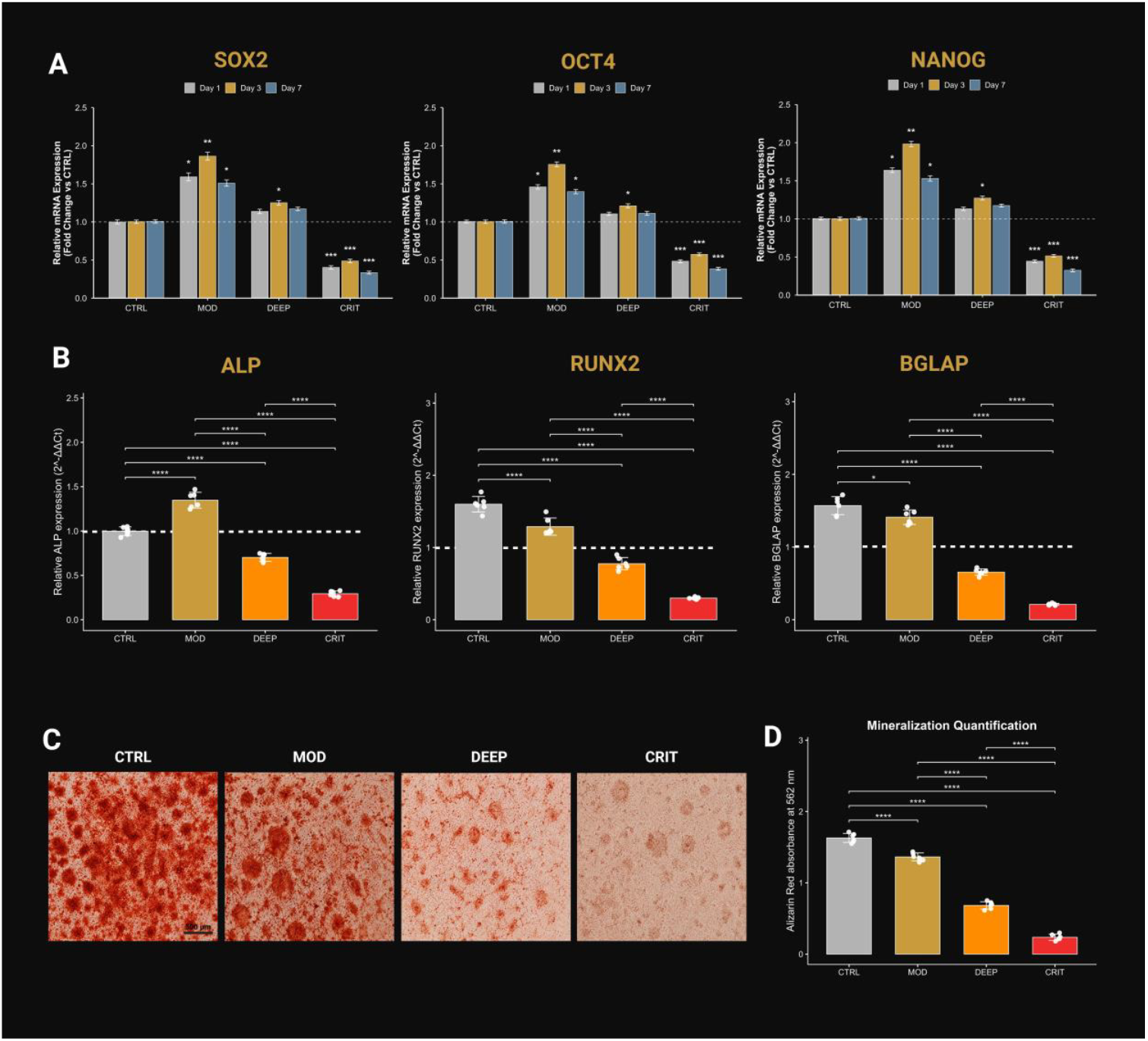
Stemness preservation and osteogenic differentiation capacity of human dental pulp stem cells (hDPSCs) following exposure to simulated subterranean hypoxic environments. A) Relative mRNA expression of SOX2, OCT4 and NANOG normalized to GAPDH and expressed as fold change relative to normoxic controls. Moderate hypoxia (10% O_2_) induced transient upregulation of stemness-associated markers, particularly at Day 3, whereas severe hypoxia (1% O_2_) significantly suppressed stemness marker expression across all evaluated timepoints. B) Relative gene expression of osteogenic markers alkaline phosphatase (ALP), runt-related transcription factor 2 (RUNX2), and bone gamma-carboxyglutamate protein (BGLAP/osteocalcin) after osteogenic induction under different oxygen tensions: normoxia (CTRL, 21% O_2_), moderate hypoxia (MOD, 10% O_2_), deep hypoxia (DEEP, 5% O_2_), and severe hypoxia (CRIT, 1% O_2_). Gene expression levels were normalized to GAPDH and expressed relative to CTRL. C) Representative micrographs of Alizarin Red S staining at Day 14 showing extracellular calcium deposition and mineralized nodule formation under the different hypoxic conditions. D) Quantification of mineralization by cetylpyridinium chloride extraction assay following Alizarin Red S staining, expressed as absorbance values at 562 nm. Increasing hypoxic severity resulted in progressive suppression of osteogenic differentiation and mineralization capacity, particularly in the CRIT group. Data are presented as mean ± standard deviation (SD) (n = 6). Statistical analysis was performed using one-way ANOVA followed by Tukey’s post hoc test. *p < 0.05, **p < 0.01, ***p < 0.001. Scale bar = 500 μm.

### Osteogenic Differentiation

Hypoxic preconditioning significantly influenced osteogenic differentiation (**Figure 4B-D**). Cells exposed to moderate hypoxia demonstrated ALP and RUNX2 expression levels comparable to or slightly higher than CTRL during early differentiation stages. In contrast, DEEP and CRIT groups showed significant reductions in ALP, RUNX2, and osteocalcin (OCN) expression following prolonged hypoxic exposure (p < 0.01), with the greatest suppression observed in CRIT cultures.

Alizarin Red staining further demonstrated reduced mineralized matrix deposition under severe hypoxia. CTRL and MOD cultures exhibited extensive mineralized nodules after 14 days of osteogenic induction, whereas DEEP cultures showed reduced staining intensity and CRIT cultures demonstrated sparse mineralization with significantly lower quantitative absorbance values compared with CTRL and MOD groups (p < 0.001).

### Overall Response to Hypoxic Modulation

Collectively, the results demonstrated a biphasic biological response of hDPSCs to oxygen deprivation. Moderate hypoxia (10% O_2_) appeared compatible with cellular adaptation and transient stemness preservation without substantial oxidative or apoptotic damage. Conversely, prolonged severe hypoxia (1% O_2_) induced progressive oxidative stress accumulation, mitochondrial dysfunction, apoptosis, and impaired osteogenic differentiation.

## Discussion

The present study demonstrated that oxygen tension critically modulates hDPSC viability, oxidative stress response, mitochondrial function, stemness maintenance, apoptosis, and osteogenic differentiation in a concentration-dependent manner. A biphasic biological response to hypoxia was observed: moderate oxygen reduction (10% O_2_) appeared compatible with cellular adaptation and transient stemness preservation, whereas prolonged severe hypoxia (1% O_2_) induced progressive cellular dysfunction and loss of regenerative potential [18,19,29].

One of the main findings was that moderate hypoxia preserved metabolic activity and transiently increased expression of stemness-related genes including SOX2, OCT4, and NANOG [19]. These observations support previous evidence suggesting that conventional atmospheric culture conditions may not accurately reproduce the physiological microenvironment of mesenchymal stem cells [30]. Although 21% O_2_ is routinely used as the standard in vitro condition, most stem cell niches exist under substantially lower oxygen tensions in vivo. Due to its confined vascular architecture, the dental pulp likely represents a relatively hypoxic physiological environment compared with atmospheric conditions.

The improved stemness preservation observed under moderate hypoxia may therefore reflect adaptation to more physiologically relevant metabolic conditions. Similar findings have been reported in mesenchymal stem cells, where controlled hypoxia has been associated with enhanced clonogenicity, delayed senescence, and maintenance of undifferentiated phenotypes through HIF-mediated metabolic adaptation [31-33]. In the present study, moderate hypoxia was also associated with relatively stable ROS levels and preserved mitochondrial membrane integrity, suggesting that controlled oxygen reduction did not induce substantial oxidative stress.

Conversely, severe hypoxia exceeded the adaptive capacity of hDPSCs. Cells cultured under 1% O_2_ demonstrated reduced metabolic activity, mitochondrial depolarization, increased ROS accumulation, elevated apoptotic populations, and impaired osteogenic differentiation. These findings indicate that the beneficial effects of physiological hypoxia are highly threshold-dependent and that excessive oxygen deprivation ultimately results in pathological cellular stress [18,29].

Interestingly, severe hypoxia induced increased ROS production despite reduced oxygen availability. This phenomenon has previously been associated with mitochondrial dysfunction under critically low oxygen conditions, where impaired electron transport chain activity promotes electron leakage and oxidative damage [34]. The marked mitochondrial depolarization observed in the CRIT group further supports the role of mitochondrial dysfunction in severe hypoxia-mediated injury. The coexistence of oxidative stress, mitochondrial dysfunction, and apoptosis additionally suggests an interconnected degenerative response induced by prolonged oxygen deprivation [19].

Another important finding was the suppression of osteogenic differentiation following prolonged severe hypoxia. Osteogenic markers including ALP, RUNX2, and osteocalcin were significantly downregulated in severely hypoxic groups, accompanied by reduced extracellular mineralization. Osteogenic differentiation is an energy-demanding process requiring active extracellular matrix synthesis and mitochondrial activity; therefore, severe oxygen deprivation may impair osteogenic commitment through both metabolic limitation and oxidative-stress-related signaling alterations [29]. In contrast, moderate hypoxia did not significantly impair osteogenic differentiation and appeared compatible with early osteogenic marker expression.

Interestingly, moderate hypoxia did not significantly impair osteogenic differentiation and in some early stages appeared compatible with osteogenic marker expression. This observation further supports the concept that moderate oxygen reduction may preserve stem-cell functionality while avoiding the detrimental effects associated with severe hypoxia [18,29,31,32]. Such findings may hold translational relevance for regenerative dentistry and tissue engineering strategies, particularly regarding the optimization of stem cell expansion protocols and biomaterial culture environments [33].

Several limitations should be acknowledged. The study was conducted entirely in vitro using commercially available hDPSCs and therefore cannot fully reproduce the complexity of the in vivo pulpal microenvironment [18]. Additionally, oxygen tension in vivo is influenced by dynamic vascular perfusion, inflammation, and nutrient availability, which were not replicated in the present model [30]. The study also focused primarily on short-term hypoxic exposure, while longer-term fluctuating oxygen conditions may produce different biological responses. Finally, molecular pathways including HIF-1α signaling and autophagy were not directly investigated and warrant further evaluation.

Future studies should investigate dynamic oxygen cycling models, inflammatory co-stimulation, and three-dimensional culture systems capable of reproducing more physiologically relevant oxygen gradients relevant to regenerative endodontics [5,6].

## Acknowledgements

This research received no external funding. The authors acknowledge the technical support provided by the Department of Clinical Sciences and Stomatology at Università Politecnica delle Marche for providing computational and statistical analysis resources. The authors are grateful to Dr. Vittorio Collamati (Occupational physician, private practicioner) for his guidance regarding occupational environmental models and atmospheric parameter standardization. The authors declare no conflict of interest related to this study. The authors further declare that no industrial interests were involved in the study design, data interpretation, manuscript preparation, or decision to publish.

## Data Availability Statement

The datasets generated and analyzed during the current study are available from the corresponding author upon reasonable request.

### Declaration of generative AI and AI-assisted technologies in the manuscript preparation process

During the preparation of this work the authors used Microsoft Copilot in order to ease the readability of the manuscript. After using this tool, the authors reviewed and edited the content as needed and take full responsibility for the content of the published article.

## Notes

### Competing Interest Statement

The authors have declared no competing interest.

### Summary of Updates

There was a typo in the title which has been corrected in this new version

